# Validity of freehand three-dimensional ultrasound system in measurement of three-dimensional surface shape of shoulder muscles

**DOI:** 10.1101/2021.11.18.468912

**Authors:** Jun Umehara, Norio Fukuda, Shoji Konda, Masaya Hirashima

## Abstract

Accurate measurement of muscle morphology is crucial for assessing skeletal muscle capacity. Although the freehand three-dimensional ultrasound (3DUS) system is a promising technique for assessing muscle morphology, its accuracy has been validated mainly in terms of volume by examining lower limb muscles. The purpose of this study was to validate 3DUS in the measurements of 3D surface shape and volume by comparing them with MRI measurements while ensuring the reproducibility of participant posture by focusing on the shoulder muscles. The supraspinatus, infraspinatus, and posterior deltoid muscles of 10 healthy males were scanned using 3DUS and MRI while secured by an immobilization support customized for each participant. A 3D surface model of each muscle was created from the 3DUS and MRI methods, and the agreement between them was assessed. For the muscle volume, the mean difference between the two models was within 0.51 cm^3^ for all muscles. For the surface shape, the distances between the closest points of the two models was calculated for every point on the 3DUS surface model. The results showed that the median (third quartile) of the distances was less than 1.21 mm (1.89 mm) for all muscles. These results suggest that, given the above error is permitted, 3DUS can be used as an alternative to MRI in measuring volume and surface shape, even for the shoulder muscles.

## 1. Introduction

The capacity of a skeletal muscle depends on its morphology, including its volume and length (Fukunaga et al., 2001; Lieber and Ward, 2011; Narici et al., 2016). Thus, accurate measurements of muscle morphology in vivo are important for musculoskeletal modelling and assessment of changes due to aging or training. Magnetic resonance imaging (MRI) is considered the “gold standard” for direct measurement of muscle morphology in vivo (Holzbaur et al., 2007; Mitsiopoulos et al., 1998). However, MRI is expensive, time-consuming, not always available, and is sometimes restrictive for a variety of postures due to the narrowness of the gantry. As a relatively inexpensive and flexible alternative, freehand three-dimensional ultrasound (3DUS) was proposed (Mozaffari and Lee, 2017; Prager et al., 2010). This technique combines two-dimensional (2D) B-mode ultrasound scanning and three-dimensional (3D) motion analysis to create 3D ultrasound images by mapping consecutive 2D images manually scanned with a probe into 3D space.

As it is cost- and time-effective and can be utilized without posture restrictions, 3DUS is promising if its accuracy is comparable to that of MRI. To date, several studies have validated the accuracy of 3DUS. First, it was validated using human cadaveric muscle (Delcker et al., 1999) or dog isolated muscle (Weller et al., 2007) by comparing the volume measurement from 3DUS with that from the water displacement method. Barber et al. (2009) was the first to validate the method using whole human muscle in vivo, showing that the mean difference between 3DUS and MRI was only 1.9 mL in volume and 3.0 mm in length measurements. Since then, its accuracy has been repeatedly demonstrated by comparing morphological measures from 3DUS with those from MRI (Barber et al., 2019) or dissection (Haberfehlner et al., 2016; Weide et al., 2017).

However, previous studies have mainly focused on volume measurements. No study has validated the measurement of the 3D surface shape of muscle, which indicates not only representative morphological quantities (i.e., volume and length) but also 3D geometric paths; these can be used to determine the moment arm of the muscle, which is another critical parameter in musculoskeletal modelling. The validation of 3D surface shape measurement would have been difficult in previous studies because the limb posture was not strictly controlled between the 3DUS and MRI measurements. This study aimed to validate the measurement of 3D surface shape while ensuring reproducibility of the posture between 3DUS and MRI measurements. In this study, rigid immobilization support was customized for each participant by molding their body shape with polyurethane foam, ensuring that the participants had the same posture for both measurements.

Another challenge of this study was to validate 3DUS by scanning the shoulder muscles, including the posterior deltoid, supraspinatus, and infraspinatus muscles. Previous studies mainly focused on the lower limb muscles, such as the triceps surae, semitendinosus, and vastus lateralis, the paths of which are relatively straight. To capture the deltoid curvature accurately, the probe must be moved along the curvature by dramatically changing its orientation. Thus, if the calibration between the probe and laboratory coordinates is not sufficiently accurate, the reconstruction of the 3D image should be collapsed. As the supraspinatus is a deep muscle surrounded by bones, proper depth setting in the ultrasound system and scan technique are necessary to clearly visualize it on the ultrasound image. Furthermore, multiple sweeps are necessary to capture the infraspinatus due to its broadness, increasing the importance of calibration accuracy.

In summary, the purpose of this study was to validate 3DUS measurements of 3D surface shape as well as volume while ensuring the reproducibility of participant posture between 3DUS and MRI by focusing on the shoulder muscles, which require accurate scanning and calibration of the 3DUS system.

## 2. Methods

### 2.1. Participants

Ten healthy males (age 25.4 ± 4.9 yrs, height 173.3 ± 4.5 cm, body mass 65.3 ± 9.0 kg) participated in this study. None of the participants had a history of shoulder injury or pain in their upper limbs. The experiment was approved by the ethics committee of the National Institute of Information and Communications Technology and was conducted in accordance with the Declaration of Helsinki. All participants provided written informed consent before participating in the experiment.

### 2.2. Experimental setup

Freehand 3DUS and MRI measurements were performed on different days. The intervals between the measurements were 7.8 ± 6.9 days. Body mass was measured using a body composition analyzer (MC-980A plus, Tanita, Tokyo, Japan) on each day, confirming that the change in body mass across days was within 0.3 ± 0.7 kg. To ensure the reproducibility of the participant posture in the two measurements, a customized immobilization support was created for each participant prior to the first measurement. While participants lied prone in a comfortable posture in a rectangular box, polyurethane foam (IFF03 InstaFoam, CDR Systems, Alberta, Canada) was poured to fill the vacancy between the box and body so that the upper trunk, left upper arm, and left forearm were immobilized. The participants were asked not to move until the foam hardened, which took approximately 15 min.

### 2.3. MRI

Each participant laid prone on their immobilization support while they were scanned using a 3.0 Tesla MRI system (MAGNETOM Vida, Siemens Healthineers, Erlangen, Germany) (Figure 1A). Images of the left shoulder region were acquired from a 12-channel flexible body coil using the chemical shift-based two-point Dixon method (Dixon, 1984). The pulse sequence was 3D gradient echo with the following parameters: resolution = 0.7 × 0.7 × 1.0 mm, field of view = 350 × 350 mm, echo times (TE/TE2) = 2.2/3.8 ms, repetition time (TR) = 5.87 ms, flip angle = 5°, no. slices = 207, and interslice gap = 0 mm. Water and fat images were produced from in-phase and opposed-phase images.

**Figure 1.**
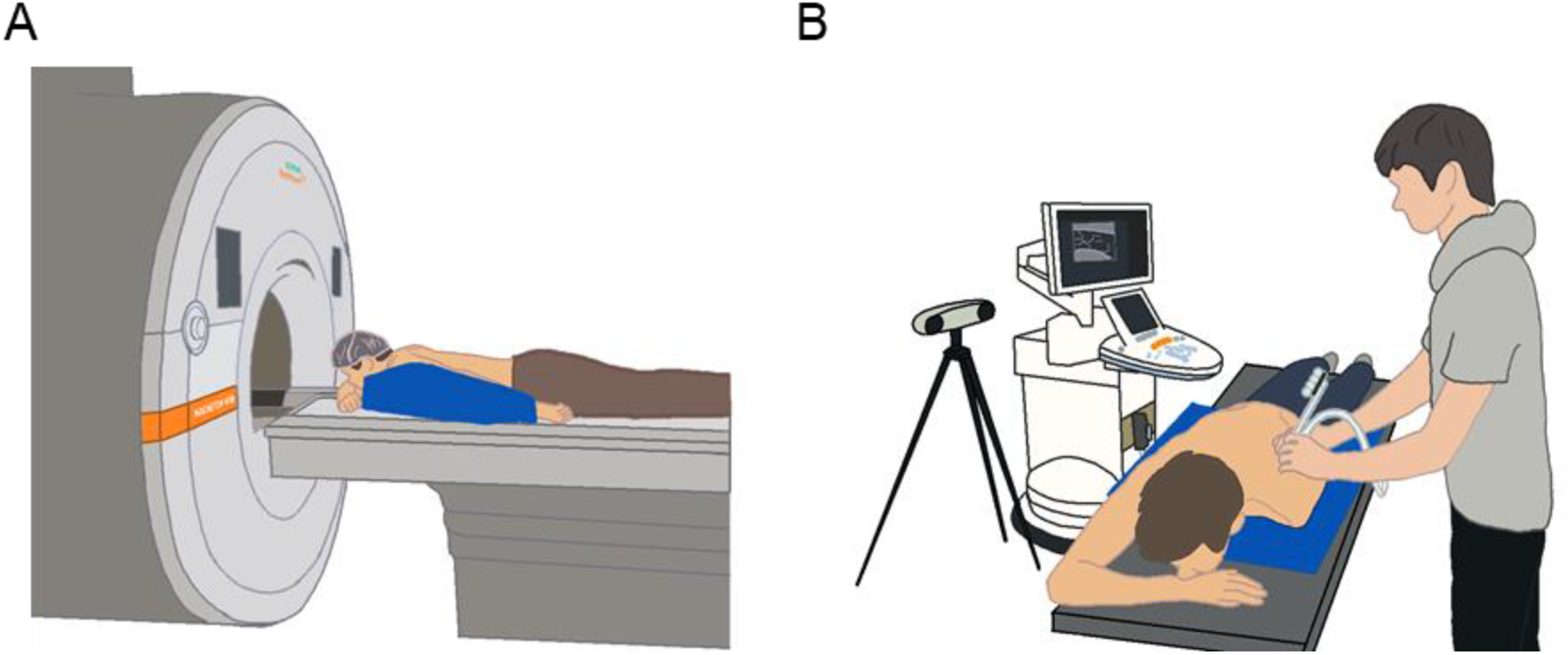
Experimental setups of MRI (A) and 3DUS (B). The participant lay prone on the rigid immobilized support customized for each participant, which is indicated blue object in this figure. The measurements of MRI and 3DUS were conducted while ensuring the reproducibility of the participant posture.

The surface shapes of the supraspinatus, infraspinatus, and whole deltoid were manually segmented by investigating the water images (Figure 2A), and a 3D surface model of each muscle was obtained (Figure 3A) using 3D Slicer (version 4.11, Harvard University, Boston, USA) (Fedorov et al., 2012). The scapula was also segmented for visualization, registration, and determination of the analysis region of the muscle volume. The manual segmentation was performed by one individual (JU) throughout the study.

**Figure 2.**
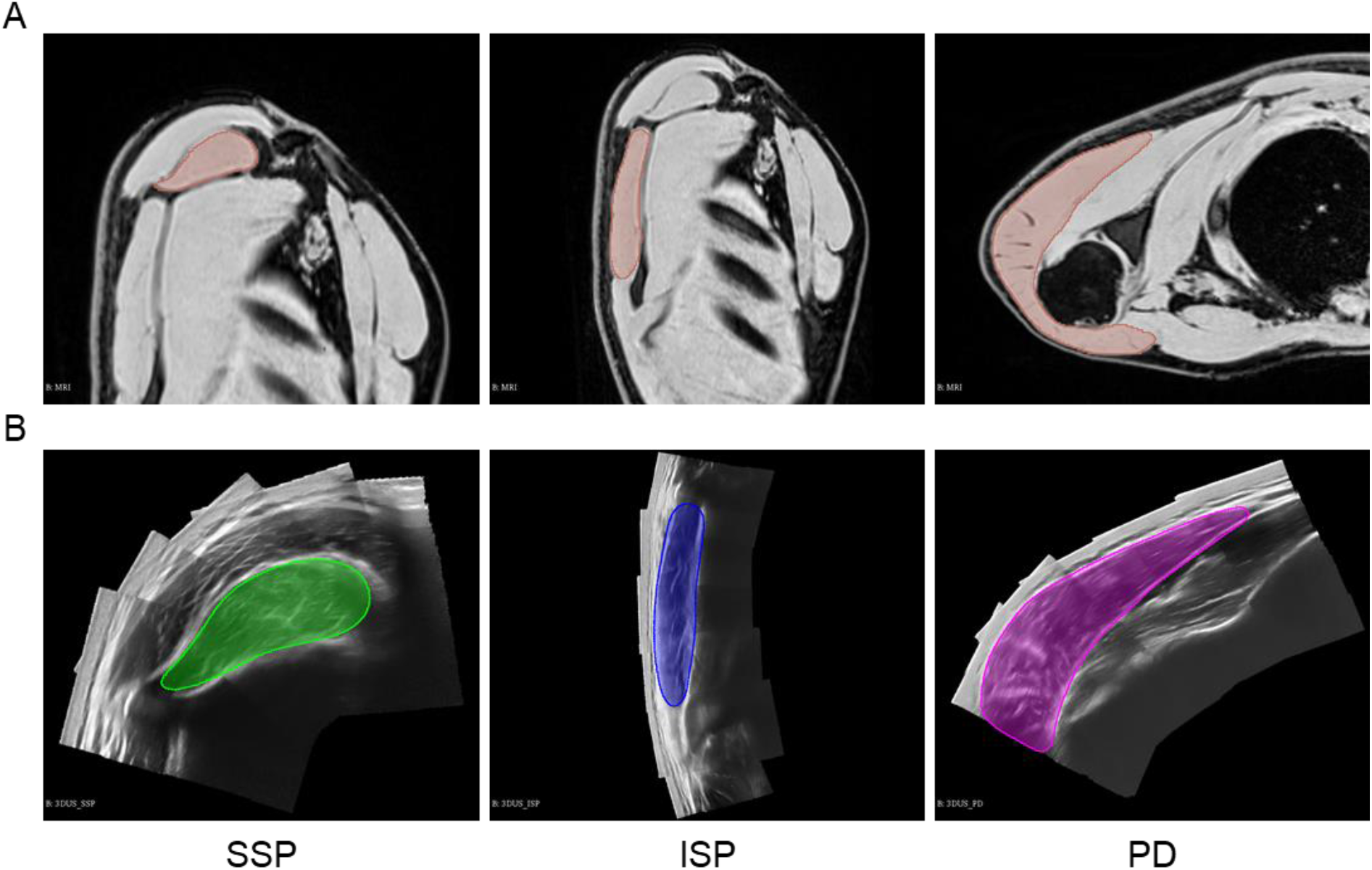
(A) MRI images. The images were sagittal plane for the supraspinatus (left) and infraspinatus (middle) and axial plane for the posterior deltoid (right). Manual segmentation was performed by depicting the muscle boundary mainly on the water images, which is represented as the colored area on each image. **(B) 3DUS images reconstructed from multiple 2D ultrasound images.** All images were measured in transverse plane along with muscle paths. The supraspinatus (left), the infraspinatus (middle), and the posterior deltoid (right) were manually segmented by delineating the muscle boundary, which is represented as the colored area on each image.

**Figure 3.**
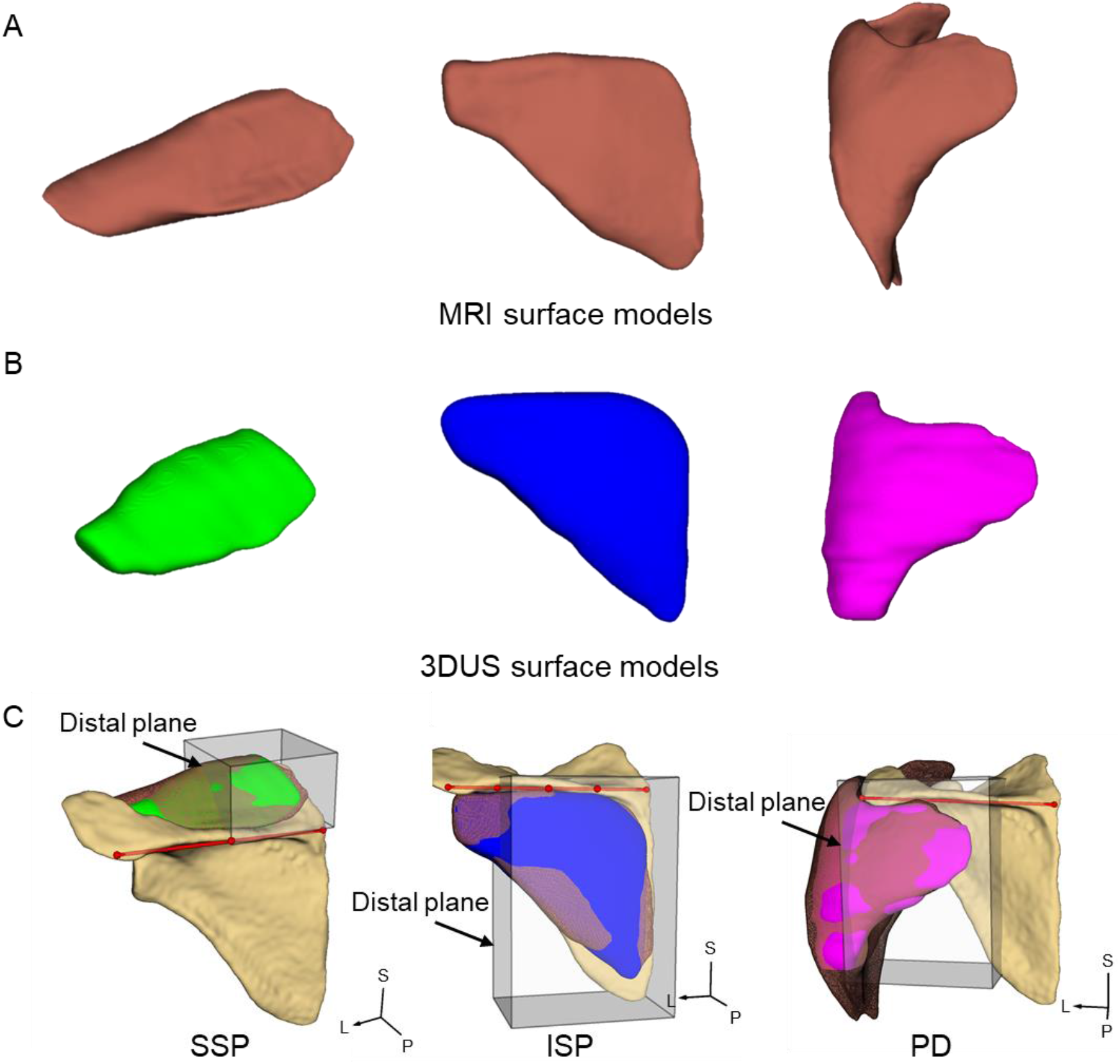
Surface shape models of MRI (A) and 3DUS (B). The surface shape model consists of point cloud and triangulation polygons. The left, middle, and right models were representative models of the supraspinatus, infraspinatus, and posterior deltoid. **(C) Surface shape models of 3DUS on MRI coordinates.** The surface shape model of 3DUS (e.g., the colored surface model) was aligned with that obtained from MRI (e.g., the wireframe model) based on the transformation matrix. The analysis region of muscle volume is indicated as a rectangular box over the 3D surface models. The red line and dots show the axis of the scapular spine and each point used for defining the analysis region. SSP, supraspinatus muscle; ISP, infraspinatus muscle; PD, posterior deltoid muscle; S, superior direction; P, posterior direction; L, lateral direction.

### 2.4. Freehand 3DUS setup

2D ultrasound images were acquired using a conventional 2D ultrasound system (Aixplorer v12.3.1, Supersonic Imagine, Aix-en Provence, France) coupled with a 2–10 MHz linear transducer (footprint: 38 mm). Either 5, 6, or 7 cm was used for the depth setting in the ultrasound system depending on the location of the muscle. The 3D position and orientation of the ultrasound probe were recorded by tacking four reflective makers rigidly attached to the probe using a motion capture system (Polaris Vicra, Northern Digital Inc., Ontario, Canada). The 2D ultrasound images and 3D motion data were collected by the 3D Slicer software.

Spatial calibration was performed using a cross-wire phantom consisting of two wires crossing each other in water. We recorded as many as 50 static 2D images capturing the cross-point at various locations in the 2D image while synchronously recording the 3D position and orientation of the probe. A least-squares fit was performed to estimate the best 4 × 4 coordinate transformation matrix from the probe coordinates to laboratory coordinates. Calibration was performed separately for each depth setting. The calibration errors were 1.81, 1.98, and 2.49 mm for the of 5, 6, and 7 cm depth settings, respectively.

The accuracy of the 3D reconstruction after calibration was assessed using a half-sphere phantom (φ = 50 mm). We manually scanned the half sphere with three sweeps that were almost parallel and overlapped with adjacent sweeps while consecutively recording 2D images and 3D motion of the probe with a sampling frequency of 10 Hz. Based on the calibration matrix, the obtained 2D images were mapped into the 3D voxel array (resolution: 0.2 × 0.2 × 0.2 mm) in the laboratory coordinates. Specifically, the grayscale value of each pixel in the 2D image was allocated to a corresponding voxel. When multiple pixels were assigned to the same voxel due to the overlap between multiple sweeps, their averaged value was set for the voxel. The surface of the spherical region was manually segmented and compared with the true sphere of 50 mm diameter. The surface distance was 0.21 ± 0.14, 0.17 ± 0.11, and 0.31 ± 0.25 mm for the of 5, 6, and 7 cm depth settings, respectively.

### 2.5. Freehand 3DUS measurement

A stack of 2D ultrasound images in the transverse plane of the supraspinatus, infraspinatus, and posterior deltoid muscles was acquired by manually moving the ultrasound probe on the left shoulder region while participants laid prone on their body support (Figure 1B). The bony landmarks of the scapula and boundaries of each muscle were marked using a pen prior to the ultrasound scan. The ultrasound probe was moved along with the muscle path at a constant velocity (approximately 1 cm/s) from origin to insertion; this is referred to as “sweep”. During the scan, copious acoustic gel was used to avoid muscle deformation that could be caused by pressure from the probe. In addition, the participants were instructed not to breathe deeply during scanning to avoid respiration artifacts.

Each muscle was scanned in a separate session. First, the muscle was pre-scanned, and the acquisition parameters of the ultrasound system were optimized for each muscle to easily visualize the muscle boundary. Depth settings were selected as 5, 6, or 7 cm depending on the muscle location. Then, the whole muscle was scanned with several sweeps at a sampling frequency of 10 Hz. The number of sweeps was three to four for the supraspinatus, five to seven for the infraspinatus, and three to six sweeps for the posterior deltoid. It took 1.7 ± 0.5, 2.6 ± 0.7, and 1.6 ± 0.7 min (including sweep intervals) to stack 347 ± 45, 488 ± 184, and 344 ± 103 images for the supraspinatus, infraspinatus, and posterior deltoid muscles, respectively. The 3D image of each muscle was reconstructed based on the calibration for the depth setting used for the muscle. Each entire muscle scan was performed twice.

### 2.6. Manual segmentation on 3DUS image

Manual segmentation was performed on the first scanned image. The surface shapes of the shoulder muscles were segmented in a similarly to how the MRI (Figure 2B) and 3D surface models of each muscle were created (Figure 3B) using 3D Slicer. For the supraspinatus, the muscle region where the bones do not appear was segmented because the distal part was not clearly imaged due to the presence of the scapular spine and clavicular on the muscle. The deltoid was segmented to obtain as much of the muscle as possible without focusing the border between the posterior and middle parts.

### 2.7. Registration

To compare the muscle models of 3DUS and MRI in the same coordinates, registration was performed based on the muscle surface. The muscle surfaces of 3DUS and MRI were used to estimate a linear transformation matrix from 3DUS coordinates to MRI coordinates using the iterative closest point algorithm, which minimizes the square error between two surface models. Based on the transformation matrix, the 3DUS surface model was aligned with the MRI surface model (Figure 3C).

### 2.8. Analysis region

Muscle volume analysis was performed on the rectangular region indicated in Figure 3C. For the supraspinatus, the distal plane of the analysis region was defined as the plane that includes a half-point of the axis of the scapular spine (from the trigonum spinae to the acromion process) and is perpendicular to the axis. For the infraspinatus and posterior deltoid, the distal plane was located on the three-quarter point of the scapular spine from proximal and the acromion process, respectively. The superior-inferior and anterior-posterior planes of the analysis region were arbitrarily defined to include the muscle boundaries.

### 2.9. Validity analysis

Agreement between the 3DUS and MRI was assessed in two ways. First, Bland-Altman analysis (Bland and Altman, 1986, 1999) was used to assess the agreement in the volume measurement by testing fixed and proportional biases. Second, surface distance analysis was used to assess the agreement in the measurement of surface shape. Specifically, the surface models obtained from 3DUS and MRI were expressed using point clouds. The 3DUS surface models were represented as 163766 ± 23208 points for the supraspinatus, 430263 ± 38408 points for the infraspinatus, and 296261 ± 89051 points for the deltoid. The MRI surface models consisted of 27416 ± 3937 points for the supraspinatus, 53124 ± 4066 points for the infraspinatus, and 118680 ± 14972 points for the deltoid. For each point of the 3DUS surface model, the distance to the closest point in the MRI surface model was calculated, and then the median and third quartile of the surface distances were calculated for each muscle of each participant.

### 2.10. Reliability analysis

To assess the reliability of manual segmentation independently of the reliability of the 3DUS scan, the first scanned image was re-segmented by the same individual (JU). Agreement between the two surface models from the first scanned image with different manual segmentations was assessed in terms of volume and surface shape using intra-class correlation coefficient (ICC) and surface distance analysis, respectively.

To assess the reliability of the 3DUS scan independently of the reliability of manual segmentation, the surface model for the second scanned image was obtained based on the same manual segmentation performed on the first scanned image. Specifically, a non-rigid deformation field from the first to the second scanned image for the same muscle was obtained by image-to-image registration, and the surface model manually segmented from the first scanned image was deformed by the field to obtain the surface model for the second scanned image. Agreement between the two surface models from the first and second scanned images was assessed in terms of volume and surface shape. Reliability analysis was performed using data from five randomly selected participants.

## 3. Results

### 3.1. Validity

The individual volumes of each muscle are presented in Figure 4A. The volume derived from 3DUS was similar to that obtained using MRI. Bland-Altman analysis found that the mean differences between the 3DUS and MRI volumes were −0.24 cm^3^ for the supraspinatus, −0.16 cm^3^ for the infraspinatus, and −0.51 cm^3^ for the posterior deltoid. The 95% limits of agreement (LoA) were 2.68 cm^3^ for the supraspinatus, 5.59 cm^3^ for the infraspinatus, and 3.24 cm^3^ for the posterior deltoid (Figure 4B and Table 1). A t-test showed that the mean difference in volume was sufficiently close to zero in the supraspinatus (t = −0.544, *P* = 0.600), infraspinatus (t = −0.177, *P* = 0.864), and posterior deltoid muscles (t = −0.975, *P* = 0.355), indicating that there was no fixed bias between the two methods. Linear regression analyses revealed that the slope of the regression line of the mean volume of the difference was sufficiently close to zero for the supraspinatus (slope = −0.065, *P* = 0.282), infraspinatus (slope = 0.026, *P* = 0.579), and posterior deltoid muscles (slope = −0.012, *P* = 0.851), indicating that there was no proportional bias between the two methods. Thus, no systematic bias was found in the volume measurements between 3DUS and MRI for all muscles.

**Figure 4.**
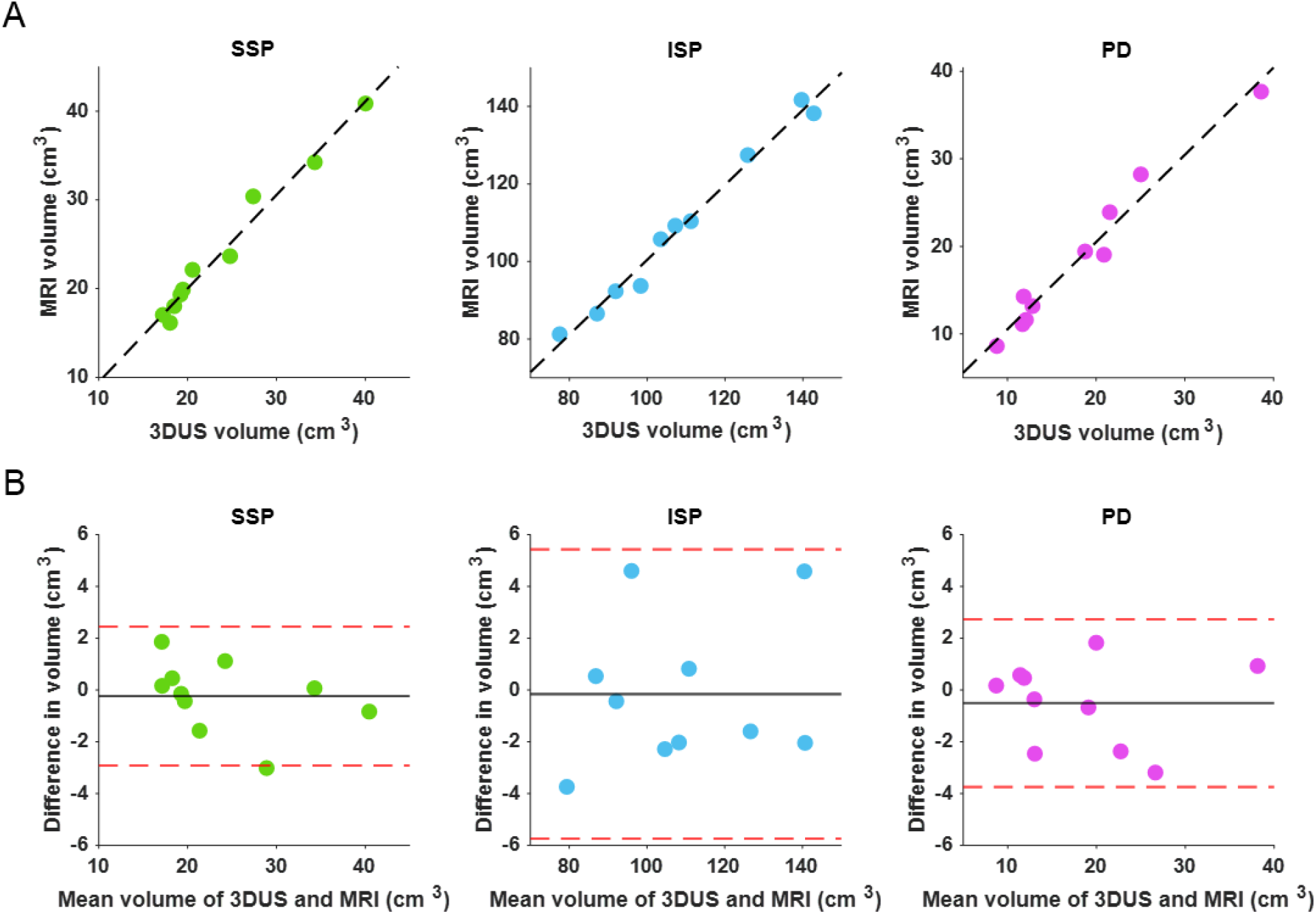
(A) Individual muscle volume measured by MRI and 3DUS. Each dot indicates individual participants. The dash line represents the linear regression line. Regression equation and coefficient of determinations (R2) were estimated using the least-squares method as follows: SSP, MRI volume = 1.05*(3DUS volume) - 1.03 (R2 = 0.98); ISP, MRI volume = 0.97*(3DUS volume) + 3.88 (R2 = 0.98); PD, MRI volume = 1.00*(3DUS volume) + 0.59 (R2 = 0.97). **(B) Bland-Altman plot for muscle volume measurement. Each dot indicates individual participants.** The black solid line represents the mean difference between the 3DUS and MRI. The red dash lines indicate upper and lower 95% LoAs. The upper and lower 95% LoAs were −2.92 and 2.45 for the SSP, −5.74 and 5.43 for the ISP, and −3.75 and 2.73 for the PD. SSP, supraspinatus muscle, ISP, infraspinatus muscle; PD, posterior deltoid muscle; LoA, limits of agreement.

**Table 1.** Agreement of muscle volume between 3DUS and MRI.

| Muscle | 3DUS (cm<sup>3</sup>) | MRI (cm<sup>3</sup>) | Mean difference (cm<sup>3</sup>) | 95% LoA (cm<sup>3</sup>) |
| --- | --- | --- | --- | --- |
| SSP | 23.94 ± 7.77 | 24.18 ± 8.28 | -0.24 ± 1.37 | 2.68 |
| ISP | 108.49 ± 21.81 | 108.65 ± 21.24 | -0.16 ± 2.85 | 5.59 |
| PD | 18.20 ± 8.94 | 18.71 ± 9.05 | -0.51 ± 1.65 | 3.24 |

Data are presented as mean and standard deviation across participants. SSP, supraspinatus muscle; ISP, infraspinatus muscle; PD, posterior deltoid muscle; 3DUS, three-dimensional ultrasound; MRI, magnetic resonance imaging; LoA, limits of agreement.

The results of the surface distance analysis are shown in Figure 5 and Table 2. The box plot in Figure 5A indicates the distribution of the closest point distances between the 3DUS and MRI surface models for each participant. Figure 5B shows the distribution of surface distances pooled across all participants. The median and third quartile of the distances were 0.71 mm and 1.21 mm for the supraspinatus, 0.94 mm and 1.66 mm for the infraspinatus, and 1.12 mm and 1.89 mm for the posterior deltoid, respectively.

**Figure 5.**
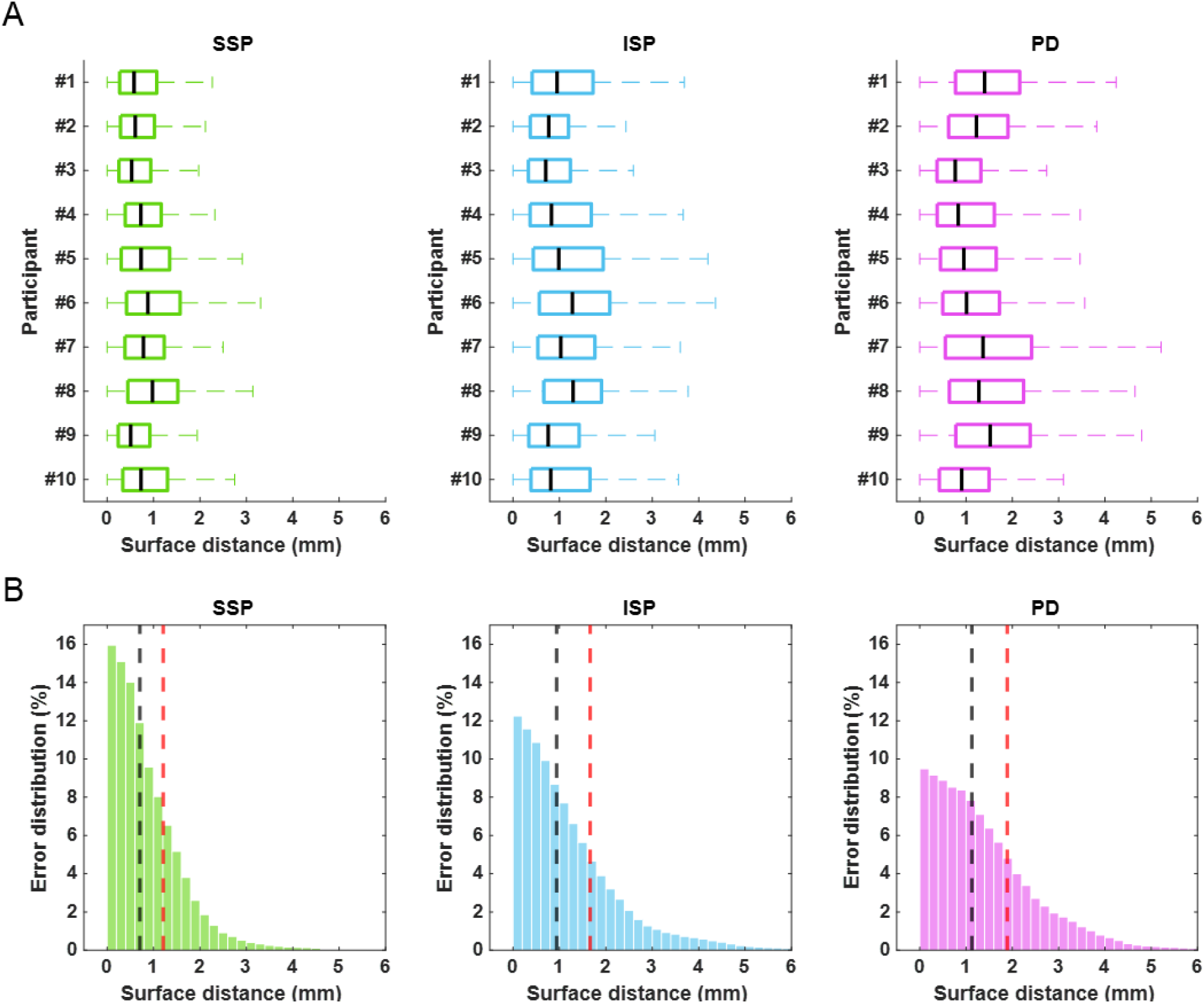
(A) Individual surface distances between 3DUS and MRI. Each box plot represents individual participants (from #1 to #10). The black vertical line within box indicates median value. The whiskers indicate maximum and minimum value. Outliers are not displayed in this figure for clear visualization. **(B) Distribution of Surface distances between 3DUS and MRI for all participants.** Each histogram pools the surface distance data of all participants. The black and red dash lines indicate the median and third quartile of the surface distance between the 3DUS and MRI. The median and third quartile are calculated by averaging them across participants. SSP, supraspinatus muscle, ISP, infraspinatus muscle; PD, posterior deltoid muscle.

**Table 2.** Surface distance between the 3DUS and MRI.

| Muscle | Median (mm) | Third quartile (mm) |
| --- | --- | --- |
| SSP | $0.71 \pm 0.15$ | $1.21 \pm 0.23$ |
| ISP | $0.94 \pm 0.21$ | $1.66 \pm 0.30$ |
| PD | $1.12 \pm 0.27$ | $1.89 \pm 0.39$ |

Data are presented as mean and standard deviation across participants. SSP, supraspinatus muscle; ISP, infraspinatus muscle; PD, posterior deltoid muscle.

### 3.2. Reliability

The reliability of the manual segmentation is described in Table 3. The ICCs for the volume measurement between the two surface models from the first scanned image with different manual segmentations were greater than 0.97 for all muscles. The surface distance analysis showed that the medians and third quartiles of the distances were less than 1.05 mm and 1.57 mm for all muscles, respectively.

**Table 3.** Reliability of the 3DUS manual segmentation.

| Muscle | Muscle volume |  | Surface distance |  |
| --- | --- | --- | --- | --- |
|  | Mean difference (cm<sup>3</sup>) | ICC | Median (mm) | Third quartile (mm) |
| SSP | -0.64 ± 1.04 | 0.99 | 0.45 ± 0.08 | 0.80 ± 0.13 |
| ISP | -0.73 ± 2.74 | 0.97 | 1.05 ± 0.49 | 1.57 ± 0.60 |
| PD | -0.42 ± 0.98 | 0.99 | 0.67 ± 0.06 | 1.20 ± 0.14 |

Data are presented as mean and standard deviation across participants. SSP, supraspinatus muscle; ISP, infraspinatus muscle; PD, posterior deltoid muscle; ICC, intraclass correlation coefficient.

The reliability of the scanning is described in Table 4. The ICCs for the volume measurement between the surface models from the first and second scanned images were greater than 0.91 for all muscles. The surface distance analysis showed that the medians and third quartiles of the distances were less than 0.47 mm and 0.81 mm for all muscles, respectively.

**Table 4.** Reliability of the 3DUS scanning.

| Muscle | Muscle volume |  | Surface distance |  |
| --- | --- | --- | --- | --- |
|  | Mean difference (cm<sup>3</sup>) | ICC | Median (mm) | Third quartile (mm) |
| SSP | -0.77 ± 1.66 | 0.96 | 0.38 ± 0.44 | 0.51 ± 0.53 |
| ISP | -0.54 ± 4.90 | 0.91 | 0.44 ± 0.12 | 0.75 ± 0.21 |
| PD | -0.73 ± 0.89 | 0.99 | 0.47 ± 0.06 | 0.81 ± 0.10 |

Data are presented as mean and standard deviation across participants. SSP, supraspinatus muscle; ISP, infraspinatus muscle; PD, posterior deltoid muscle; ICC, intraclass correlation coefficient.

## 4. Discussion

This study aimed to validate the accuracy of 3DUS for measurements of the 3D surface shape and muscle volume in shoulder muscles by comparing it with MRI measurements. To this end, we assessed the agreement between the two methods in terms of muscle volume and surface shape by scanning shoulder muscles while ensuring the reproducibility of participant posture. Our results show that there was no systematic bias between 3DUS and MRI for volume measurement. Moreover, the median of the surface distances between 3DUS and MRI was less than approximately 1 mm for the measurement of surface shape. To the best of our knowledge, this is the first study to verify the accuracy of 3DUS for assessing the morphology and 3D surface shape of shoulder muscles.

The validity of 3DUS has only previously been ensured only for lower limb muscles. Barber et al. (2009) first validated the accuracy of 3DUS in the measurement of the whole muscle volume in the medial gastrocnemius, with a mean difference of 1.9 mL and 95% LoA of 18 mL between 3DUS and MRI. Later, Noorkoiv et al. (2019) examined the validity of 3DUS by measuring the medial gastrocnemius of typical developing children and children with cerebral palsy and reported that the mean difference between 3DUS and MRI was 0.14 cm^3^, and the 95% LoA ranged from - 4.10 to 4.39 cm^3^. Other previous studies also reported similar levels of accuracy of 3DUS in the rectus femoris of living humans in comparison with MRI (e.g., mean difference and 95% LoA were 0.53 cm^3^ and 2.14 cm^3^, respectively) (MacGillivray et al., 2009) and in the semitendinosus of cadavers in comparison with the dissection method (e.g., mean difference and 95% LoA were 0.0 cm^3^ and 4.6 cm^3^, respectively) (Haberfehlner et al., 2016); in these cases, only a portion of muscle was used for the comparisons. Considering the accuracy of previous studies, our results for muscle volume are acceptable for all muscles.

Notably, this study was the first to determine whether 3DUS can be used to measure the 3D surface shape of muscles. To compare the 3D surface shape, we rigorously reproduced the participant posture between 3DUS and MRI by rigid immobilization support customized for each participant. Consequently, the medians and even third quartiles of surface distance between 3DUS and MRI were less than 1.12 and 1.89 mm for all muscles, respectively. This measurement error is thought to be sufficiently small when considering the measurement difference in muscle volume, which is attributed to the surface distances. Our results indicate that 3DUS is a promising technique to assess not only the muscle morphology but also 3D surface shape of muscles, elucidating the 3D geometric path, which eventually determines the moment arm of the muscle.

Another challenge of this study was to validate the measurement accuracy of 3DUS for shoulder muscles. No study has previously validated the 3DUS measurement of shoulder muscles because shoulder muscles are thought to be difficult to accurately obtain ultrasound images of due to their anatomical features, such as curvature, deepness, and broadness. Moreover, sufficiently constructing 3DUS images from images scanned by multiple sweeps requires an accurate calibration from the probe coordinates to the laboratory ones, making accurate assessment by 3DUS more difficult. In fact, one previous study described that misalignment occurred in the reconstruction process of 3DUS images from two sweeps (Barber et al., 2019). In the present study, we could accurately scan and construct a 3DUS image of the shoulder muscles from multiple sweeps. Thus, this study deserves attention in terms of the methodology of 3DUS, including accurate scanning and calibration.

The reliabilities of manual segmentation and scanning were assessed in terms of muscle volume and surface shape. The ICCs for the volume measurement were greater than 0.91, and the median of the surface distance was less than approximately 1 mm in both the manual segmentation and scanning for all muscles. Previous studies reported that an ICC of more than 0.9 is considered reliable for repeated manual segmentation and ultrasound scans (Barber et al., 2019; Barber et al., 2009; MacGillivray et al., 2009). Therefore, the reliability of the proposed methodology was verified.

In conclusion, this study confirmed the agreement between 3DUS and MRI for measuring the muscle volume and 3D surface shape between the two methods under reproducible participant posture. Our findings indicate that, given that the above-mentioned error is permitted, 3DUS can be used as an alternative to MRI in measuring volume and surface shape, even for shoulder muscles.

## Acknowledgement

This work was supported by Grant-in-Aid for JSPS Fellows (20J01660) and Grant-in-Aid for Scientific Research (B) (18H03149; 21H03320).

## Author contributions

J.U., N.F., S.K., and M.H. conceived and designed the study; J.U. and N.F. performed data acquisition and analysis; J.U., N.F., S.K., and M.H. interpreted the data; J.U. and M.H. drafted and revised the manuscript critically for important intellectual content; All authors approved final version of the manuscript.

